# Support Vector Machine based prediction models for drug repurposing and designing novel drugs for colorectal cancer

**DOI:** 10.1101/2023.10.20.563215

**Authors:** Avik Sengupta, Rahul Kumar

## Abstract

Colorectal cancer (CRC) has witnessed a concerning increase in incidence and poses a significant therapeutic challenge due to its poor prognosis. There is a pressing demand to identify novel drug therapies to combat CRC. In this study, we addressed this need by utilizing the library of CRC pharmacological profiles of anticancer drugs and developed QSAR models for prediction of alternative and promiscuous anti-cancer compounds for CRC treatment. Our QSAR models demonstrated their robustness by achieving high correlation of determination (R^2^) after 10-fold cross validation. For 12 CRC cell lines, R^2^ ranges from 0.609-0.827. Highest performance was achieved for SW1417 and GP5d cell lines with R^2^, 0.827 and 0.786, respectively. Further, we listed the most common descriptors in the drug profiles of the CRC cell lines and we also checked the correlation of the descriptors with the drug activity. The KRFP314 fingerprint was the predominantly occurring descriptor, with the KRFPC314 fingerprint following closely in prevalence within the drug profiles of the CRC cell lines. Beyond predictive modeling, we also successfully validated drug-to-oncogene relationship via *in-silico* methods and identified FDA-approved drugs which could be used as potential anti-CRC drugs, paving the way for subsequent *in vitro* or *in vivo* experiments to validate their efficacy. To provide easy accessibility and utility of our research findings, we have incorporated these models into a free-to-use user-friendly prediction webserver named, “ColoRecPred” hosted on project.iith.ac.in/cgntlab/colorecpred. This web-based tool can be used to screen for potential anti-cancer compounds for CRC.

## Introduction

Colorectal cancer (CRC) has third highest incidence and second highest mortality rate of 10% and 9.4% respectively across the globe, as per GLOBOCAN 2020[1], notably affecting both males and females[1]. Some widely used drugs approved for treating CRC are cetuximab, oxaliplatin, 5-FU, and tucatinib[2]. To combat the increasing specter of drug resistance, clinicians and researchers have banked on combinational drug therapies, such as CAPOX, FOLFIRI, FU-LV, and XELOX, among others[2]. Even though this strategic shift towards combination therapy has been useful in many CRC treatments, but other strategies need to be introduced continuously to combat CRC[3]. The mechanism of drug resistances is being studied widely; therefore, many mechanisms have been recently identified[4]. Primary mechanisms include reduced drug activation, an aberration in downstream signaling processes, drug transport aberration, and changes in drug targets[4]. This emphasizes the clinical importance of augmenting the current available drug arsenal to enhance therapeutic methodologies for CRC. Implementing machine learning based *in-silico* methods, e.g. Quantitative Structure Activity Relationship (QSAR) models, are attractive approach to bypass the time and cost exhaustive traditional drug discovery process [5]. The *in-silico* methods can be used to screen large chemical libraries to predict novel drugs for CRC and boost drug discovery and development[5]. There is a continuous effort to improve the drug arsenal for CRC worldwide. Recently, many new targets have been used for drug development using 3D-QSAR models. The recently published drug targets are Interleukin-6, and DNA Topoisomerase II, where 3D-QSAR models have been developed for CRC to find drugs that show anticancer activity[6,7]. Likewise, in this study, we have developed QSAR models for 12 CRC cell lines to identify putative drugs for CRC. The QSAR models’ development is done based on the high-throughput pharmacological data available from Genomics of Drug Sensitivity in Cancer (GDSC). These models can be utilized for drug repurposing of FDA approved against CRC. Moreover, they can also hasten screening of large chemical libraries to identify novel CRC drugs[8]. Our model will be useful for the research fraternity to complement the ongoing research to identify novel drug candidates for CRC, which can be taken further for experimental validations. For the community-wide utilization of QSAR models developed in this study, we have integrated these models in a webserver called “ColoRecPred”, which is freely available at project.iith.ac.in/cgntlab/colorecpred. Figure 1 describes the overall study design adopted in this study.

**Figure 1.**
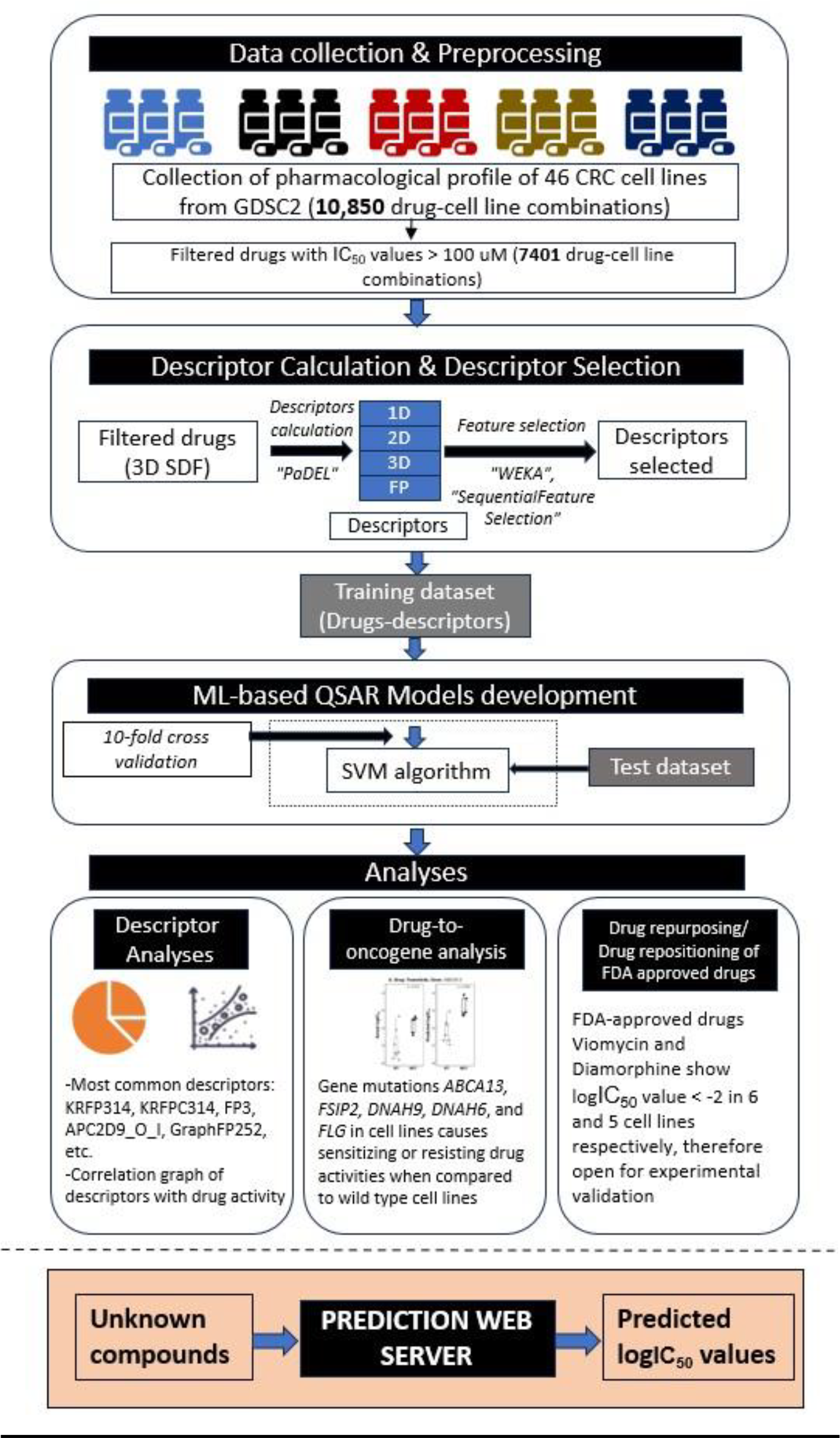
The overall study design for the development of QSAR models and prediction webserver. In this study, we have procured pharmacological drug data from GDSC for 46 cell lines of CRC, then fed the drugs to PaDEL software for descriptor calculation. To reduce the number of descriptors, we performed feature selection using WEKA tool and f-stepping process. The pharmacological drug data and descriptors were used as training and test datasets for developing QSAR models for 46 cell lines, out of which 12 cell lines showed coefficient of determination of > 0.6. Using these QSAR models, we performed descriptor analyses, drug-to-oncogene relationship analyses (for validation of our developed models) and drug repurposing/drug repositioning analyses. Finally, we hosted our developed QSAR models on a free prediction webserver to enable prediction of drug activities of unknown compounds.

## Methods

### Pharmacological Data

For this study, we have downloaded the pharmacological screens against CRC cell lines from the Genomics of Drug Sensitivity in Cancer (GDSC) database (https://www.cancerrxgene.org). A dataset of 297 anticancer drugs and their respective natural logarithmic IC_50_ values were obtained across 46 CRC cell lines. We had total of 10,850 drug-cell lines combinations each with IC_50_ value and then applied cut-off of 100μM to remove inactive drugs, reducing the total drug-cell lines combinations to 7401. The total number of drugs for all 46 cell lines are shown in Supplementary Figure 1. We extracted the individual cell line screened data with their respective logIC_50_ values. PubChem compound IDs (CIDs) of drugs were also retained to obtain the chemical structures of the drugs.

### Chemical Structure of Drugs

The chemical structures of all the above drugs were downloaded in Spatial Data File (SDF) format from PubChem using their CIDs (https://pubchem.ncbi.nlm.nih.gov). These structures were in 2D format; therefore, they were subjected to 3D conversion using the RDKit toolkit in python[9] followed by energy minimization using Merck Molecular Force Field 1994 (MMFF94) force field [10–14] in OpenBabel software (version 3.1.1)[15].

### Descriptors Calculation and Selection

We used PaDEL software[16] to calculate the 1875 chemical descriptors (1D, 2D, and 3D) across 75 descriptor types like number of atoms count, topological, bond count, atom count, 3D autocorrelation, moment of inertia, RDF, WHIM, etc. and 12 different types of binary fingerprints like FP, ExtFP, GraphFP, SubFP, SubFPC, etc. Figure 2 shows the distribution of the descriptor types. Total descriptor counts equal to 18066, across 75 descriptor types. For descriptor selection, we implemented “RemoveUseless” function (removes descriptors with no variation or very high variation)[19] to preprocess the dataset, and then implemented “CfssubsetEval” Attribute Evaluator (evaluates the predictive ability of a descriptor and intercorrelation among other descriptors)[17,19] and “BestFirst” Ranker (evaluates features based on Greedy Hillclimbing and Backtracking mechanism)[17] in WEKA to select the descriptors.

**Figure 2.**
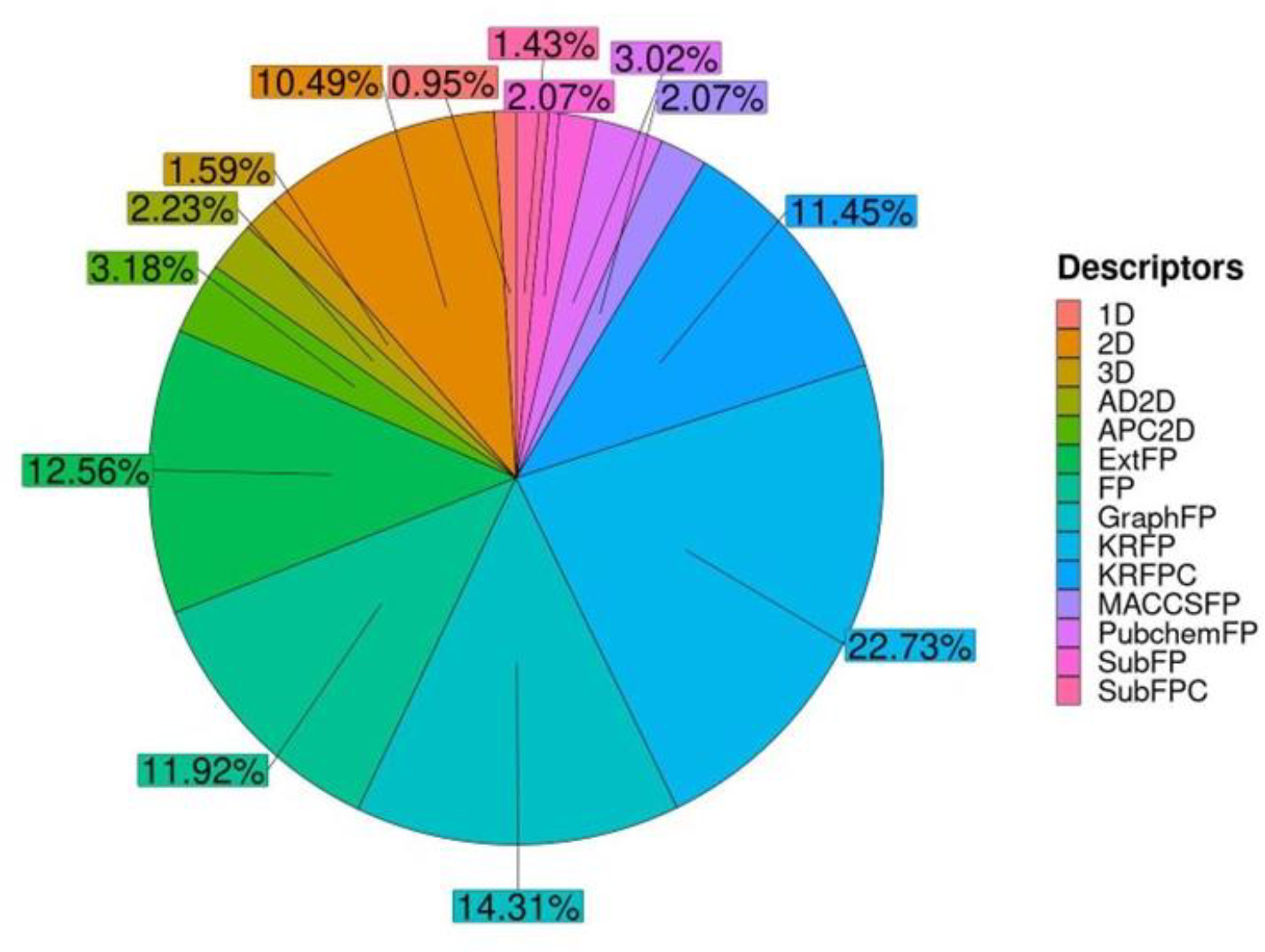
Graphical representation of the overview of the distribution of the descriptor types across the 12 selected cell lines. Pie chart representing the types of descriptors (1D, 2D, 3D and Binary Fingerprints) across the 12 cell lines.

### QSAR Model development

To develop QSAR models, we used Support Vector Machine (SVM) algorithm using “scikit-learn” (version 1.2)[20] library in Python. We implemented 10-fold cross validation to avoid over-fitting and assessed the model performances using various statistical indices i.e., Pearson’s correlation coefficient (R), coefficient of determination (R^2^), mean squared error (MSE), mean average error (MAE) and Root mean square error (RMSE). To identify the robust QSAR models, we used cut-off of R^2^ > 0.6. Selected models were further subjected to “F-stepping” to reduce the number of descriptors using “SequentialFeatureSelection” function in “Mlxtend” library in Python environment[21]. During model development, we maintained the drugs to descriptor ratio for each of the selected cell lines close to 2:1 or greater to reduce the chances of overfitting[22].

### Drug-to-Oncogene Relation

We used QSAR models to recapitulate drug-to-oncogene relationships in CRC. We downloaded mutation data of CRC cell lines from the COSMIC Cell Line Project database (v97, https://cancer.sanger.ac.uk/cell_lines)[23]. From the COSMIC mutation data, we removed mutations defined as “Unknown” and “Substitution - coding silent”. We selected five genes i.e., *ABCA13, DNAH6, DNAH9, FSIP2*, and *FLG*, which were mutated in at least five CRC cell lines (Supplementary Table 1). For these five genes, we identified drugs with significant differences in their respective logIC_50_ between wild type and mutant cell lines (p < 0.05) and predicted their logIC_50_ using QSAR models of respective wild type and mutant cell lines. Then, we compared the predicted logIC_50_ of respective drugs in wild type and mutated cell lines to recapitulate the drug-to-oncogene relation obtained from the experimentally known logIC_50_.

**Table 1:** Performance Measures calculated for the 12 cell lines on training dataset.

| S. No. | Cell Line | No. of Drugs | Performance before F-stepping |  |  |  |  | Performance after F-stepping |  |  |  |  |
| --- | --- | --- | --- | --- | --- | --- | --- | --- | --- | --- | --- | --- |
|  |  |  | No. of Desc<br>ripto<br>rs | R<sup>2</sup> | R | RMS<br>E | MAE | No. of Desc<br>ripto<br>rs | R<sup>2</sup> | R | RMS<br>E | MAE |
| 1. | COLO-678 | 56 | 50 | 0.654 | 0.827 | 0.961 | 0.735 | 29 | 0.687 | 0.845 | 0.915 | 0.695 |
| 2. | HT-115 | 138 | 95 | 0.601 | 0.777 | 1.154 | 0.887 | 68 | 0.64 | 0.804 | 1.097 | 0.834 |
| 3. | SW620 | 122 | 86 | 0.572 | 0.793 | 1.719 | 1.263 | 44 | 0.639 | 0.827 | 1.578 | 1.096 |
| 4. | SW1463 | 122 | 79 | 0.621 | 0.792 | 1.224 | 0.873 | 47 | 0.662 | 0.835 | 1.155 | 0.779 |
| 5. | COLO-205 | 142 | 67 | 0.707 | 0.85 | 1.454 | 1.161 | 46 | 0.732 | 0.868 | 1.392 | 1.052 |
| 6. | GP5d | 151 | 102 | 0.713 | 0.845 | 1.306 | 0.958 | 55 | 0.786 | 0.887 | 1.126 | 0.848 |
| 7. | HT-29 | 146 | 134 | 0.601 | 0.782 | 1.491 | 1.097 | 43 | 0.707 | 0.844 | 1.277 | 0.908 |
| 8. | KM12 | 131 | 87 | 0.689 | 0.846 | 1.484 | 1.147 | 46 | 0.749 | 0.871 | 1.334 | 0.992 |
| 9. | SW1417 | 76 | 208 | 0.693 | 0.852 | 1.087 | 0.837 | 48 | 0.827 | 0.927 | 0.814 | 0.52 |
| 10. | MDST8 | 134 | 92 | 0.668 | 0.829 | 1.634 | 1.421 | 64 | 0.71 | 0.869 | 1.527 | 1.103 |
| 11. | SK-CO-1 | 142 | 75 | 0.693 | 0.841 | 1.455 | 1.159 | 50 | 0.747 | 0.88 | 1.322 | 1.051 |
| 12. | CCK-81 | 154 | 138 | 0.56 | 0.764 | 1.311 | 0.961 | 89 | 0.609 | 0.789 | 1.236 | 0.899 |

### Drug Repurposing/ Drug Repositioning

We obtained the 1627 FDA approved drugs from DrugBank (https://go.drugbank.com/) in the SDF format. Their chemical descriptors and fingerprints were calculated using PaDEL (as described above) and we predicted their logIC_50_ values using QSAR models developed in this study. To select FDA approved drugs with putative anticancer activity, we applied a cutoff of logIC_50_ value <= -2.

## Results

### QSAR Models Performance

To evaluate the performance of QSAR models, we adopted four statistical indices, a) Pearson’s correlation coefficient (R), b) coefficient of determination (R^2^), c) Root Mean Square Error (RMSE), and d) Mean Absolute Error (MAE). We developed individual QSAR models for 46 CRC cell lines by splitting the datasets into train (80%) and test (20%). To reduce overfitting, we applied 10-fold cross validation within the training dataset across all the cell lines. After cross-validation step, we applied cutoff of R^2^ > 0.6 to select the best QSAR models. With this selection criteria, we obtained 12 cell lines (Table 1) and proceeded with them for downstream analysis. For each QSAR model, we ensured drugs to descriptors ratio of 2:1 or greater. To achieve this, we further reduced the number of descriptors using “F-stepping” as mentioned in the Methods section. The descriptor and drug numbers for the selected 12 cell lines are shown in Table 1. The performances were measured at two different descriptor counts, one after the “CfsSubsetEval” module in WEKA and the other after F-stepping. It was observed that, in most of the cell lines, performance of the models after F-stepping improved (Table 1). Highest performance was achieved for SW1417 cell lines (R^2^ = 0.827), and the lowest performance was for CCK-81 cell lines (R^2^=0.609) on training dataset (Table 1). Further, we tested QSAR models on test dataset and obtained performance ranging from 0.603 to 0.882 (Table 2). Supplementary Figure 2 shows the scatter plots (with linear fit) between actual and predicted logIC_50_ values for the 12 CRC cell lines.

**Table 2:**
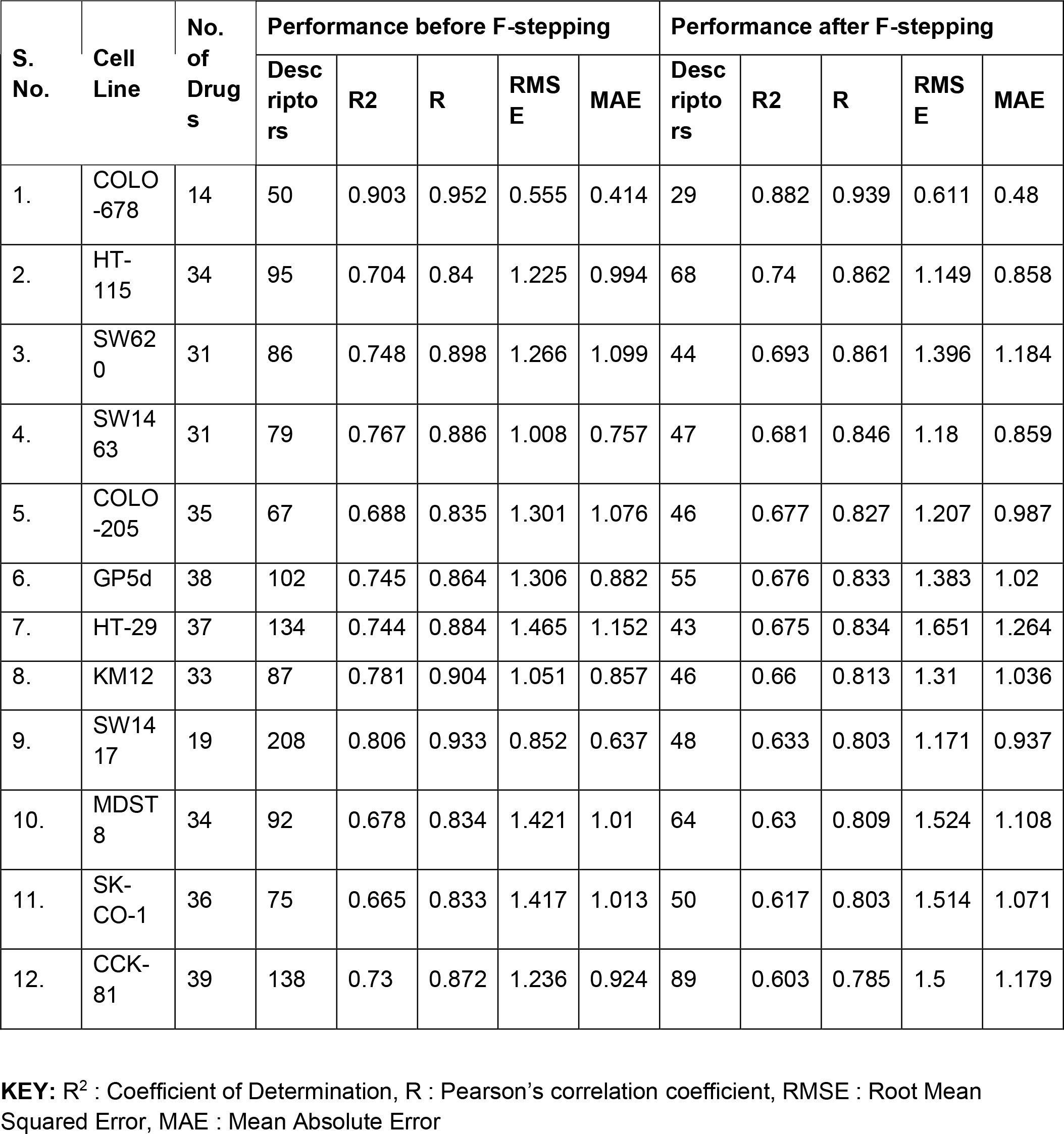
Performance Measures calculated for the 12 cell lines on test dataset.

### Descriptor Analysis

We listed the most occurring descriptors and fingerprints and, we found that fingerprints KRFP314 and KRFPC314 were the two-most occurring descriptors across 12 selected cell lines (occurring in nine and seven cell lines, respectively), as shown in Supplementary Table 2. We also checked for correlation of these frequently occurring descriptors with drug activity (logIC_50_) and observed a negative correlation between the descriptor and logIC_50_ values. This suggests better drug activity (i.e., lower logIC_50_) while a particular descriptor is present (Figure 3 and Supplementary Table 3).

**Figure 3.**
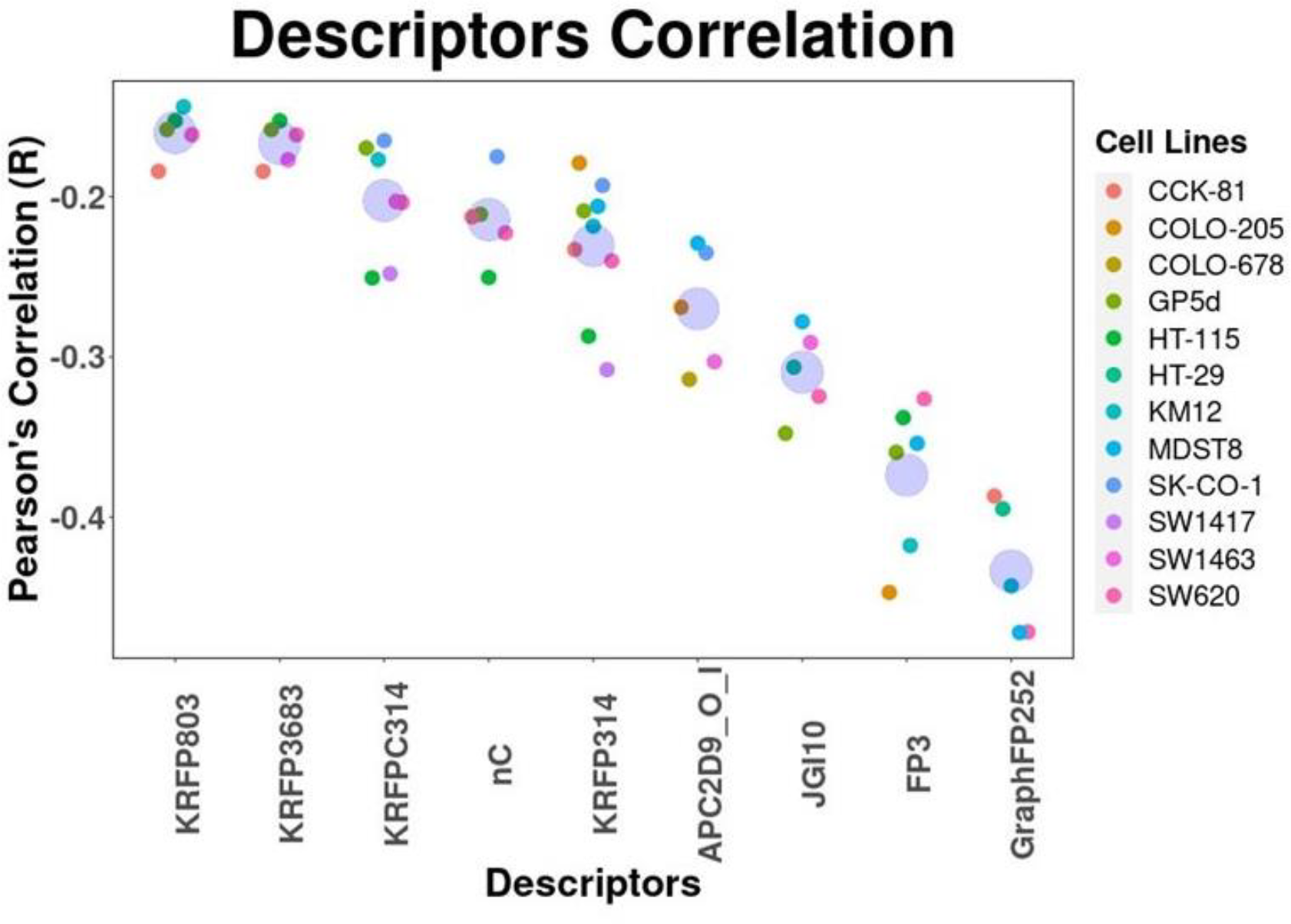
Most occurring descriptors in the drug profiles of the 12 cell lines and their correlation coefficient. Pearson’s correlation coefficient was calculated for the 9 descriptors. We selected the descriptors which were present in at least 5 cell line drug profiles.

### Drug-to-oncogene relation validation

We used QSAR models to rehash the drug-to-oncogene relationships obtained from the experimental data. We identified the association between *ABCA13* and Trametinib, where *ABCA13* mutated cell lines were less sensitive for Trametinib as compared to wild type cell lines (P=0.019). Using QSAR models developed in this study, we predicted the logIC_50_ of Trametinib for these cell lines and observed a similar trend with predicted logIC_50_ (P = 0.0034) (Figure 4A). We found another association between *FSIP2* and SGC0946, where *FSIP2* mutated cell lines were more sensitive to SGC0946 (P=0.0059). We predicted the logIC_50_ using QSAR models and recapitulated this association (P=0.0057) (Figure 4B). We found more such drug to oncogene relations and recapitulated them using QSAR models developed in this study (Supplementary Figure 3), which highlights the predictive power of these models.

**Figure 4.**
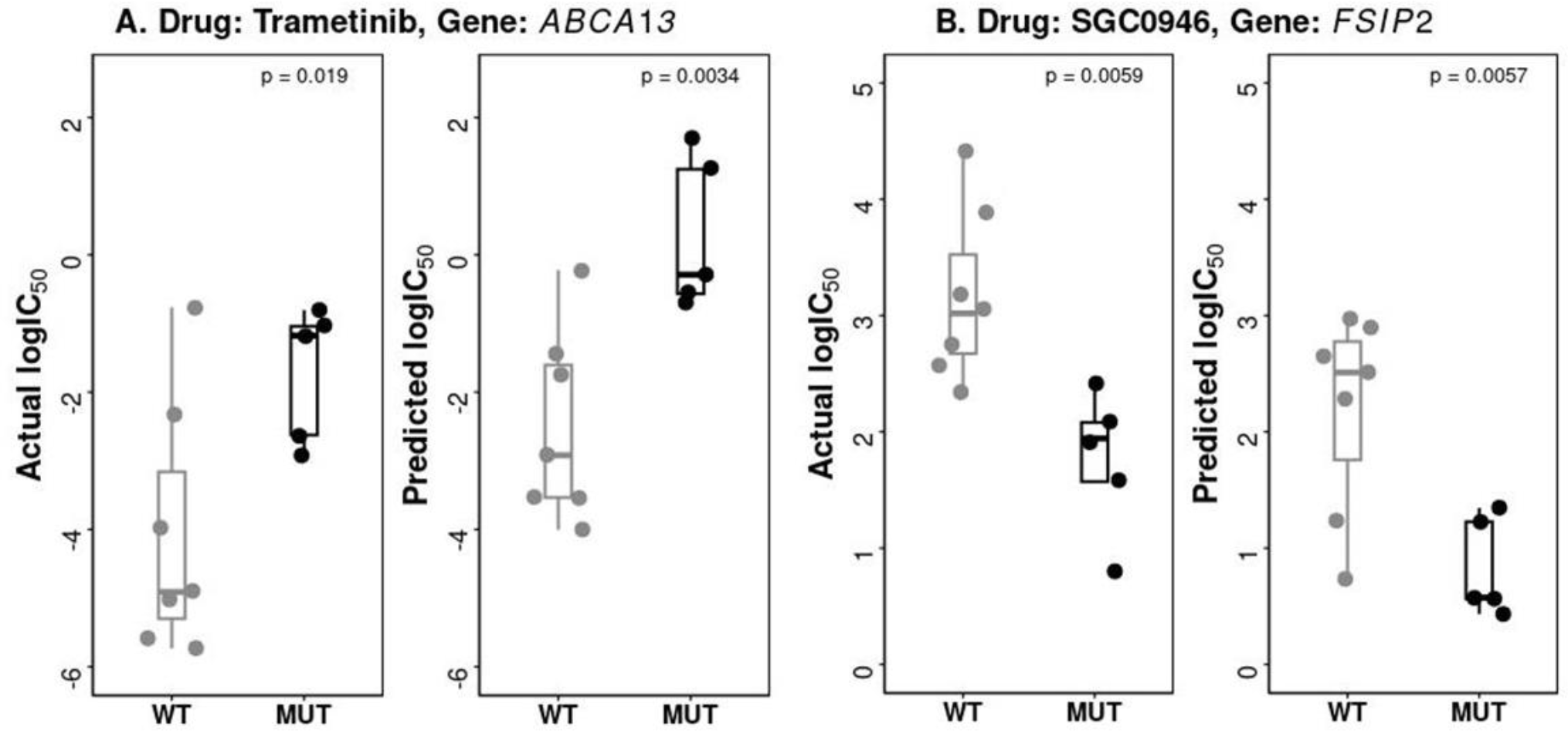
Validation of drug-to-oncogene relation using QSAR models. (A). Mutations in *ABCA13* reduced the sensitivity of CRC cell lines for Trametinib in both actual (experimental) and predicted logIC_50_ and (B). Mutations in *FSIP2* increased the sensitivity of CRC cell lines for SGC0946 in both actual (experimental) and predicted logIC_50_.

### FDA-approved drugs analysis

We further extended our QSAR models to repurpose FDA-approved drugs for CRC. We predicted the logIC_50_ of 1627 FDA approved drugs across the 12 CRC cell lines using our QSAR models. Of these, we filtered the drugs with predicted logIC_50_ =< -2 uM for each of the 12 cell lines to select the drugs with prominent activity. We found 11 drugs with logIC50 =< -2 in at least five cell lines out of 12 (Table 3). Of these 11 drugs, six were known anti-cancer drugs, three were anti-microbial that have been shown to have an antineoplastic effect[24–45], and two drugs i.e., viomycin (anti-microbial agent) and diamorphine (analgesic) have yet not been experimentally validated as anti-cancer drugs[46–48].

**Table 3:**
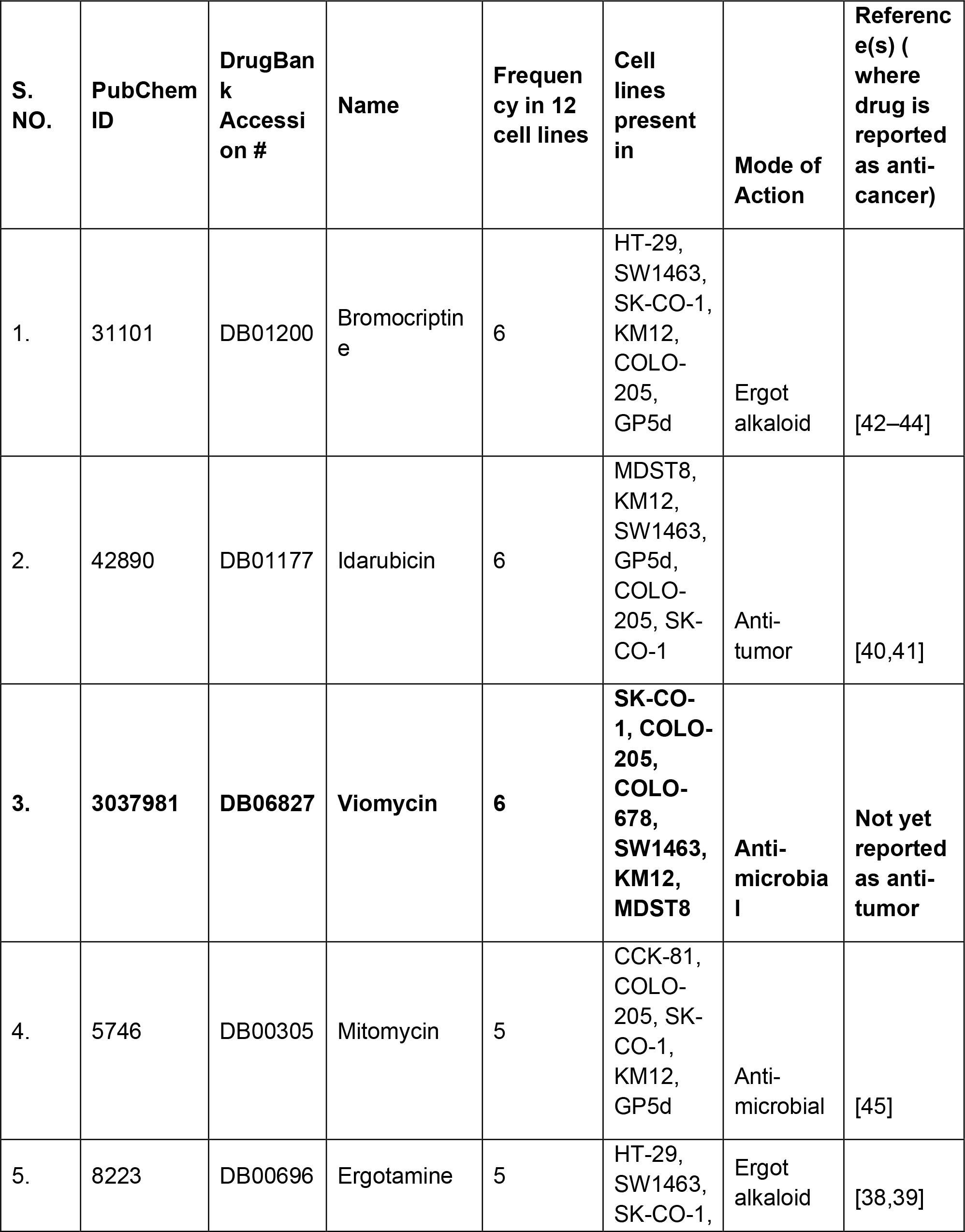
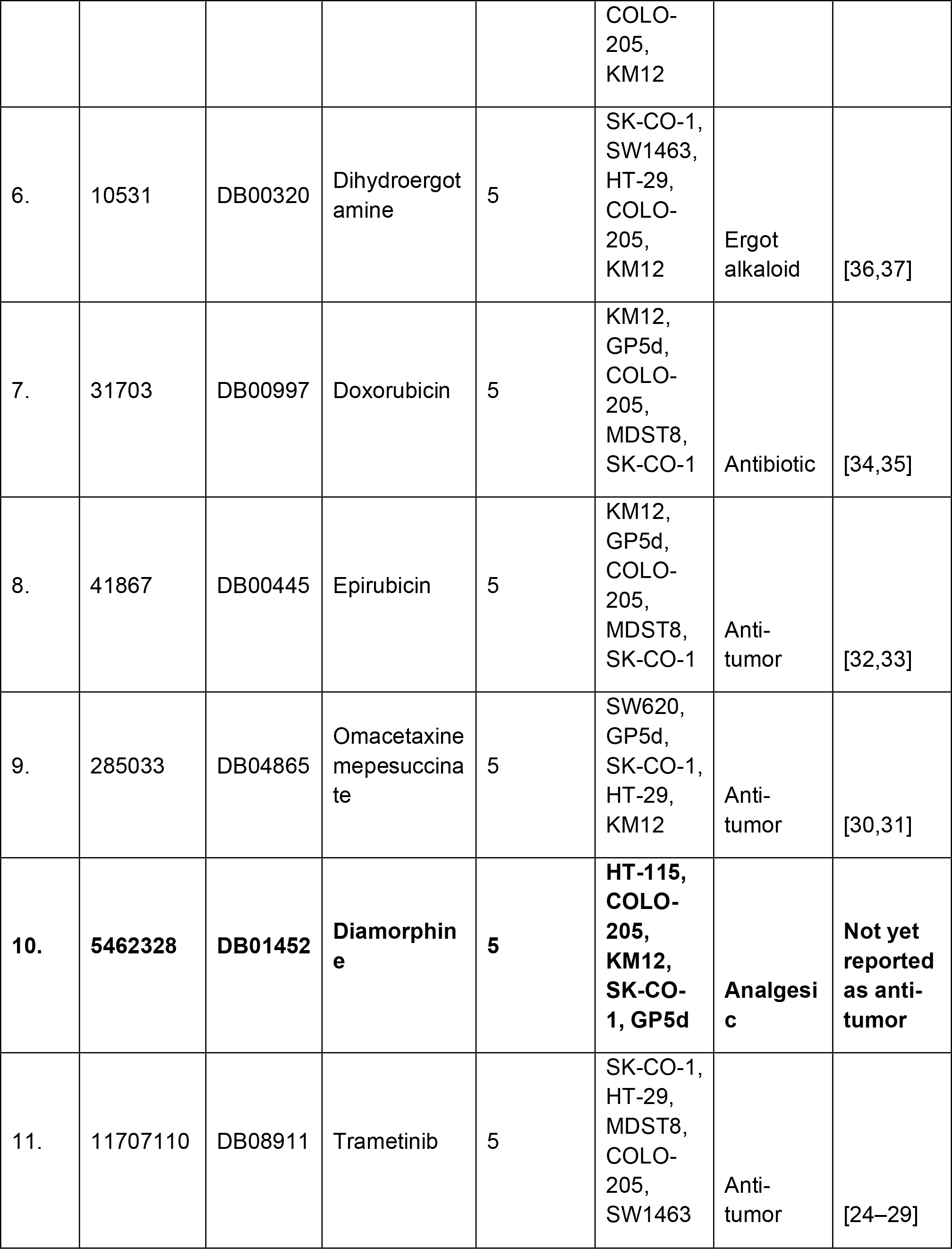
Most potent Drugs after analyzing FDA approved drugs by our QSAR models.

### Web Server

We integrated these QSAR models in a webserver, named “ColoRecPred”. The webserver shall be used to predict the anti-cancer activity (logIC_50_ values) of the unknown molecules or compounds against CRC cell lines. The “Predict” page in the webserver allows users to paste 2D structure (in SDF format) of an unknown compound and select the CRC cell lines for which the user wants to predict logIC_50_ values. On submitting the queries, the webserver shall return the logIC_50_ values of the compound for each selected cell lines in a tabular format. “ColoRecPred” is freely available at https://project.iith.ac.in/cgntlab/colorecpred.

## Discussion

In the context of CRC, there is an urgency to swiftly develop alternative therapies and medications to address the challenge of high incidence and mortality[49]. Further, there is the problem of prolonged timeline of discovery of new drugs (∼ 10 to 15 years), which ends up being very expensive and complex[50]. Computational strategies have been developed and adapted in both pharmaceutical industries and academia to boost the process of drug discovery and drug development [51]. With computational strategies in mind, we have developed robust QSAR models based on SVM. These models will facilitate the screening of potential drugs for CRC treatment, whether as standalone therapies or in combination with conventional drugs. In our study, we developed QSAR models for 12 CRC cell lines to design novel drug compounds against CRC. Drugs have symbolic codes or structures which are quantifiable, known as descriptors which can confer the potent drug activity of the molecule. We utilized the correlation of these descriptors with the drug activity to develop the QSAR models. In descriptors analyses, we found the presence of various 1D, 2D, 3D descriptors and molecular fingerprints across the drugs. With the help of descriptors selection, we could effectively design models with performance (R^2^) ranging from 0.609 to 0.827. To show the robustness of our QSAR models, we implemented them to recapitulate drug-to-oncogene relations identified using experimental data. We successfully recapitulated the associations between the genes *ABCA13, FLG, FSIP2, DNAH6* and *DNAH9* and drugs Trametinib, SGC0946, Dinaciclib, Sabutoclax, Vincristine and SCH772984 respectively. These genetic associations, if studied deeper, shall open dimensions to discover novel biomarkers or drug targets for CRC. To be further affirmative of the results, experimental validation of these drug-to-oncogene relationships is necessary. We anticipate that QSAR models developed in this study will be critical for drug repurposing and designing of novel drugs against CRC.

## Acknowledgements

We are thankful to Council of Scientific & Industrial Research (CSIR) for providing the necessary monetary support in form of stipend and contingency, and Indian Institute of Technology Hyderabad (IITH) for providing the necessary infrastructure to help us successfully conduct our research work. We acknowledge Kavita Kundal for critical of the manuscript.

## Author Contributions

Conceptualization and Design, Data acquisition, analysis and interpretation: AS and RK; Writing of the manuscript draft: AS and RK. All authors contributed to the article and approved the submitted version.

## Funding

Research seed grant (#SG/IITH/F271/2022-23/SG-109) from Indian Institute of Technology Hyderabad.

## Conflict of Interest

All authors declare that they have no conflicts of interest.

## Notes

### Competing Interest Statement

The authors have declared no competing interest.

## References

[1] H. Sung, J. Ferlay, R.L. Siegel, M. Laversanne, I. Soerjomataram, A. Jemal, F. Bray, Global Cancer Statistics 2020: GLOBOCAN Estimates of Incidence and Mortality Worldwide for 36 Cancers in 185 Countries, CA Cancer J Clin. 71 (2021) 209–249. 10.3322/CAAC.21660.

[2] Drugs Approved for Colon and Rectal Cancer - NCI, (n.d.). https://www.cancer.gov/about-cancer/treatment/drugs/colorectal (accessed April 23, 2023).

[3] K. Van Der Jeught, H.C. Xu, Y.J. Li, X. Bin Lu, G. Ji, Drug resistance and new therapies in colorectal cancer, World J Gastroenterol. 24 (2018) 3834. 10.3748/WJG.V24.I34.3834.

[4] Q. Wang, X. Shen, G. Chen, J. Du, Drug Resistance in Colorectal Cancer: From Mechanism to Clinic, Cancers (Basel). 14 (2022). 10.3390/CANCERS14122928.

[5] S. Moshawih, A.F. Lim, C. Ardianto, K.W. Goh, N. Kifli, H.P. Goh, Q. Jarrar, L.C. Ming, Target-Based Small Molecule Drug Discovery for Colorectal Cancer: A Review of Molecular Pathways and In Silico Studies, Biomolecules. 12 (2022). 10.3390/BIOM12070878.

[6] A. Rai, V. Raj, M.H. Aboumanei, A.K. Singh, A.K. Keshari, S.P. Verma, S. Saha, Pharmacophore, 3D-QSAR Models and Dynamic Simulation of 1,4-Benzothiazines for Colorectal Cancer Treatment, Comb Chem High Throughput Screen. 20 (2017). 10.2174/1386207320666170509153137.

[7] D.M. Khaled, M.E. Elshakre, M.A. Noamaan, H. Butt, M.M. Abdel Fattah, D.A. Gaber, A Computational QSAR, Molecular Docking and In Vitro Cytotoxicity Study of Novel Thiouracil-Based Drugs with Anticancer Activity against Human-DNA Topoisomerase II, Int J Mol Sci. 23 (2022). 10.3390/IJMS231911799.

[8] S.C. Peter, J.K. Dhanjal, V. Malik, N. Radhakrishnan, M. Jayakanthan, D. Sundar, D. Sundar, Quantitative Structure-Activity Relationship (QSAR): Modeling Approaches to Biological Applications, Encyclopedia of Bioinformatics and Computational Biology: ABC of Bioinformatics. 1–3 (2019) 661–676. 10.1016/B978-0-12-809633-8.20197-0.

[9] RDKit: Open-source cheminformatics. https://www.rdkit.org, (2006).

[10] T.A. Halgren, Performance of MMFF94*, John Wiley & Sons, Inc, 1996. http://journals.wiley.com/jcc.

[11] T.A. Halgren, Merck molecular force field. II. MMFF94 van der Waals and electrostatic parameters for intermolecular interactions, J Comput Chem. 17 (1996) 520–552. 10.1002/(SICI)1096-987X(199604)17:5/6<520::AID-JCC2>3.0.CO;2-W.

[12] T.A. Halgren, Merck molecular force field. III. Molecular geometries and vibrational frequencies for MMFF94, J Comput Chem. 17 (1996) 553–586. 10.1002/(SICI)1096-987X(199604)17:5/6<553::AID-JCC3>3.0.CO;2-T.

[13] T.A. Halgren, R.B. Nachbar, Merck molecular force field. IV. conformational energies and geometries for MMFF94, J Comput Chem. 17 (1996) 587–615. 10.1002/(SICI)1096-987X(199604)17:5/6<587::AID-JCC4>3.0.CO;2-Q.

[14] T.A. Halgren, Merck molecular force field. V. Extension of MMFF94 using experimental data, additional computational data, and empirical rules, J Comput Chem. 17 (1996) 616–641. 10.1002/(SICI)1096-987X(199604)17:5/6<616::AID-JCC5>3.0.CO;2-X.

[15] N.M. O’Boyle, M. Banck, C.A. James, C. Morley, T. Vandermeersch, G.R. Hutchison, Open Babel: An Open chemical toolbox, J Cheminform. 3 (2011) 1–14. 10.1186/1758-2946-3-33/TABLES/2.

[16] C.W. Yap, PaDEL-descriptor: An open source software to calculate molecular descriptors and fingerprints, J Comput Chem. 32 (2011) 1466–1474. 10.1002/JCC.21707.

[17] M. Hall, E. Frank, G. Holmes, B. Pfahringer, P. Reutemann, I.H. Witten, The WEKA data mining software, ACM SIGKDD Explorations Newsletter. 11 (2009) 10–18. 10.1145/1656274.1656278.

[18] M.A.H. and I.H.W. Eibe Frank, The WEKA Workbench. Online Appendix for “Data Mining: Practical Machine Learning Tools and Techniques,” Fourth edition, 2016.

[19] M. Hall, L. Smith, Feature subset selection: a correlation based filter approach, Proceedings of International Conference on Neural Information Processing and Intelligent Information Systems. (1998) 855–858.

[20] F. Pedregosa Fabianpedregosa, V. Michel, O. Grisel Oliviergrisel, M. Blondel, P. Prettenhofer, R. Weiss, J. Vanderplas, D. Cournapeau, F. Pedregosa, G. Varoquaux, A. Gramfort, B. Thirion, O. Grisel, V. Dubourg, A. Passos, M. Brucher, M. Perrot and Édouardand, andÉdouard Duchesnay, Fré. Duchesnay EDOUARDDUCHESNAY, Scikit-learn: Machine Learning in Python, Journal of Machine Learning Research. 12 (2011) 2825–2830. http://jmlr.org/papers/v12/pedregosa11a.html (accessed April 24, 2023).

[21] S. Raschka, MLxtend: Providing machine learning and data science utilities and extensions to Python’s scientific computing stack, J Open Source Softw. 3 (2018) 638. 10.21105/JOSS.00638.

[22] R. Kumar, K. Chaudhary, D. Singla, A. Gautam, G.P.S. Raghava, Designing of promiscuous inhibitors against pancreatic cancer cell lines, Sci Rep. 4 (2014) 4668. 10.1038/srep04668.

[23] J.G. Tate, S. Bamford, H.C. Jubb, Z. Sondka, D.M. Beare, N. Bindal, H. Boutselakis, C.G. Cole, C. Creatore, E. Dawson, P. Fish, B. Harsha, C. Hathaway, S.C. Jupe, C.Y. Kok, K. Noble, L. Ponting, C.C. Ramshaw, C.E. Rye, H.E. Speedy, R. Stefancsik, S.L. Thompson, S. Wang, S. Ward, P.J. Campbell, S.A. Forbes, COSMIC: the Catalogue Of Somatic Mutations In Cancer, Nucleic Acids Res. 47 (2019) D941–D947. 10.1093/NAR/GKY1015.

[24] M. Elbadawy, Y. Sato, T. Mori, Y. Goto, K. Hayashi, M. Yamanaka, D. Azakami, T. Uchide, R. Fukushima, T. Yoshida, M. Shibutani, M. Kobayashi, Y. Shinohara, A. Abugomaa, M. Kaneda, H. Yamawaki, T. Usui, K. Sasaki, Anti-tumor effect of trametinib in bladder cancer organoid and the underlying mechanism, Cancer Biol Ther. 22 (2021) 357–371. 10.1080/15384047.2021.1919004.

[25] L. Liu, H. Shi, M.R. Bleam, V. Zhang, J. Zou, J. Jing, K.E. Bachman, M. Motwani, K.W. Orford, A. Hoos, Antitumor effects of dabrafenib, trametinib, and panitumumab as single agents and in combination in BRAF -mutant colorectal carcinoma (CRC) models., Journal of Clinical Oncology. 32 (2014) 3513–3513. 10.1200/jco.2014.32.15_suppl.3513.

[26] E.L. Bolf, T.C. Beadnell, M.M. Rose, A. D’Alessandro, T. Nemkov, K.C. Hansen, R.E. Schweppe, Dasatinib and Trametinib Promote Anti-Tumor Metabolic Activity, Cells. 12 (2023) 1374. 10.3390/cells12101374.

[27] M.Y.K. Ho, M.J. Morris, J.L. Pirhalla, J.W. Bauman, C.B. Pendry, K.W. Orford, R.A. Morrison, D.S. Cox, Trametinib, a first-in-class oral MEK inhibitor mass balance study with limited enrollment of two male subjects with advanced cancers, Xenobiotica. 44 (2014) 352–368. 10.3109/00498254.2013.831143.

[28] R. Zeiser, Trametinib, in: 2014: pp. 241–248. 10.1007/978-3-642-54490-3_15.

[29] A.K. Salama, K.B. Kim, Trametinib (GSK1120212) in the treatment of melanoma, Expert Opin Pharmacother. 14 (2013) 619–627. 10.1517/14656566.2013.770475.

[30] M. Wetzler, D. Segal, Omacetaxine as an Anticancer Therapeutic: What is Old is New Again, Curr Pharm Des. 17 (2011) 59–64. 10.2174/138161211795049778.

[31] A. Nazha, H. Kantarjian, J. Cortes, A. Quintás-Cardama, Omacetaxine mepesuccinate (synribo) – newly launched in chronic myeloid leukemia, Expert Opin Pharmacother. 14 (2013) 1977–1986. 10.1517/14656566.2013.821464.

[32] S. Tsukagoshi, [Epirubicin (4’-epi-adriamycin]., Gan To Kagaku Ryoho. 17 (1990) 151–9.

[33] R.J. Cersosimo, W.K. Hong, Epirubicin: a review of the pharmacology, clinical activity, and adverse effects of an adriamycin analogue., Journal of Clinical Oncology. 4 (1986) 425–439. 10.1200/JCO.1986.4.3.425.

[34] S. Peter, S. Alven, R.B. Maseko, B.A. Aderibigbe, Doxorubicin-Based Hybrid Compounds as Potential Anticancer Agents: A Review, Molecules. 27 (2022) 4478. 10.3390/molecules27144478.

[35] M. Kciuk, A. Gielecińska, S. Mujwar, D. Kołat, Ż. Kałuzińska-Kołat, I. Celik, R. Kontek, Doxorubicin—An Agent with Multiple Mechanisms of Anticancer Activity, Cells. 12 (2023) 659. 10.3390/cells12040659.

[36] S.H. Chang, A.Y. Lee, K.N. Yu, J. Park, K.P. Kim, M.H. Cho, Dihydroergotamine Tartrate Induces Lung Cancer Cell Death through Apoptosis and Mitophagy, Chemotherapy. 61 (2016) 304–312. 10.1159/000445044.

[37] M. He, Q. Liao, D. Liu, X. Dai, M. Shan, M. Yang, Y. Zhang, L. Zhai, L. Chen, L. Xiang, M. He, S. Li, A. Chen, L. Sun, J. Lian, Dihydroergotamine mesylate enhances the anti-tumor effect of sorafenib in liver cancer cells, Biochem Pharmacol. 211 (2023). 10.1016/J.BCP.2023.115538.

[38] A.M. Crider, C.K.L. Lu, H.G. Floss, J.M. Cassady, J.A. Clemens, Ergot alkaloids. Synthesis of nitrosourea derivatives of ergolines as potential anticancer agents, J Med Chem. 22 (1979) 32–35. 10.1021/jm00187a008.

[39] M. Mrusek, E.-J. Seo, H.J. Greten, M. Simon, T. Efferth, Identification of cellular and molecular factors determining the response of cancer cells to six ergot alkaloids, Invest New Drugs. 33 (2015) 32–44. 10.1007/s10637-014-0168-4.

[40] T. Tsuruo, T. Oh-Hara, Y. Sudo, M. Naito, Antitumor activity of idarubicin, a derivative of daunorubicin, against drug sensitive and resistant P388 leukemia., Anticancer Res. 13 (1993) 357–61. http://www.ncbi.nlm.nih.gov/pubmed/8517647 (accessed September 7, 2023).

[41] R.J. Cersosimo, Idarubicin: an anthracycline antineoplastic agent., Clin Pharm. 11 (1992) 152–67. http://www.ncbi.nlm.nih.gov/pubmed/1551297 (accessed September 7, 2023).

[42] F.M. Kamazani, F. Sotoodehnejad nematalahi, S.D. Siadat, M. Pornour, M. Sheikhpour, A success targeted nano delivery to lung cancer cells with multi-walled carbon nanotubes conjugated to bromocriptine, Scientific Reports 2021 11:1. 11 (2021) 1–15. 10.1038/s41598-021-03031-2.

[43] L. Bai, X. Li, Y. Yang, R. Zhao, E.Z. White, A. Danaher, N.J. Bowen, C. V. Hinton, N. Cook, D. Li, A.Y. Wu, M. Qui, Y. Du, H. Fu, O. Kucuk, D. Wu, Bromocriptine monotherapy overcomes prostate cancer chemoresistance in preclinical models, Transl Oncol. 34 (2023) 101707. 10.1016/J.TRANON.2023.101707.

[44] E.J. Seo, Y. Sugimoto, H.J. Greten, T. Efferth, Repurposing of bromocriptine for cancer therapy, Front Pharmacol. 9 (2018). 10.3389/FPHAR.2018.01030/FULL.

[45] M. Tomasz, Mitomycin C: small, fast and deadly (but very selective), Chem Biol. 2 (1995) 575–579. 10.1016/1074-5521(95)90120-5.

[46] M. Holm, A. Borg, M. Ehrenberg, S. Sanyal, Molecular mechanism of viomycin inhibition of peptide elongation in bacteria, Proc Natl Acad Sci U S A. 113 (2016) 978–983. 10.1073/PNAS.1517541113/-/DCSUPPLEMENTAL.

[47] S.P.S. Rana, A. Ahmed, V. Kumar, P.K. Chaudhary, D. Khurana, S. Mishra, Successful Management of a Difficult Cancer Pain Patient by Appropriate Adjuvant and Morphine Titration, Indian J Palliat Care. 17 (2011) 162. 10.4103/0973-1075.84541.

[48] H. Nersesyan, K. V. Slavin, Current aproach to cancer pain management: Availability and implications of different treatment options, Ther Clin Risk Manag. 3 (2007) 381. /pmc/articles/PMC2386360/ (accessed April 25, 2023).

[49] H. Brenner, M. Kloor, C.P. Pox, Colorectal cancer, The Lancet. 383 (2014) 1490–1502. 10.1016/S0140-6736(13)61649-9.

[50] R.C. Mohs, N.H. Greig, Drug discovery and development: Role of basic biological research, Alzheimer’s & Dementia : Translational Research & Clinical Interventions. 3 (2017) 651. 10.1016/J.TRCI.2017.10.005.

[51] A. V. Sadybekov, V. Katritch, Computational approaches streamlining drug discovery, Nature 2023 616:7958. 616 (2023) 673–685. 10.1038/s41586-023-05905-z.

