## Supplementary figures for "Support Vector Machine based prediction models for drug repurposing and designing novel drugs for colorectal cancer"

**Supplementary Figure 1:** Overview of the drugs count across all 46 cell lines

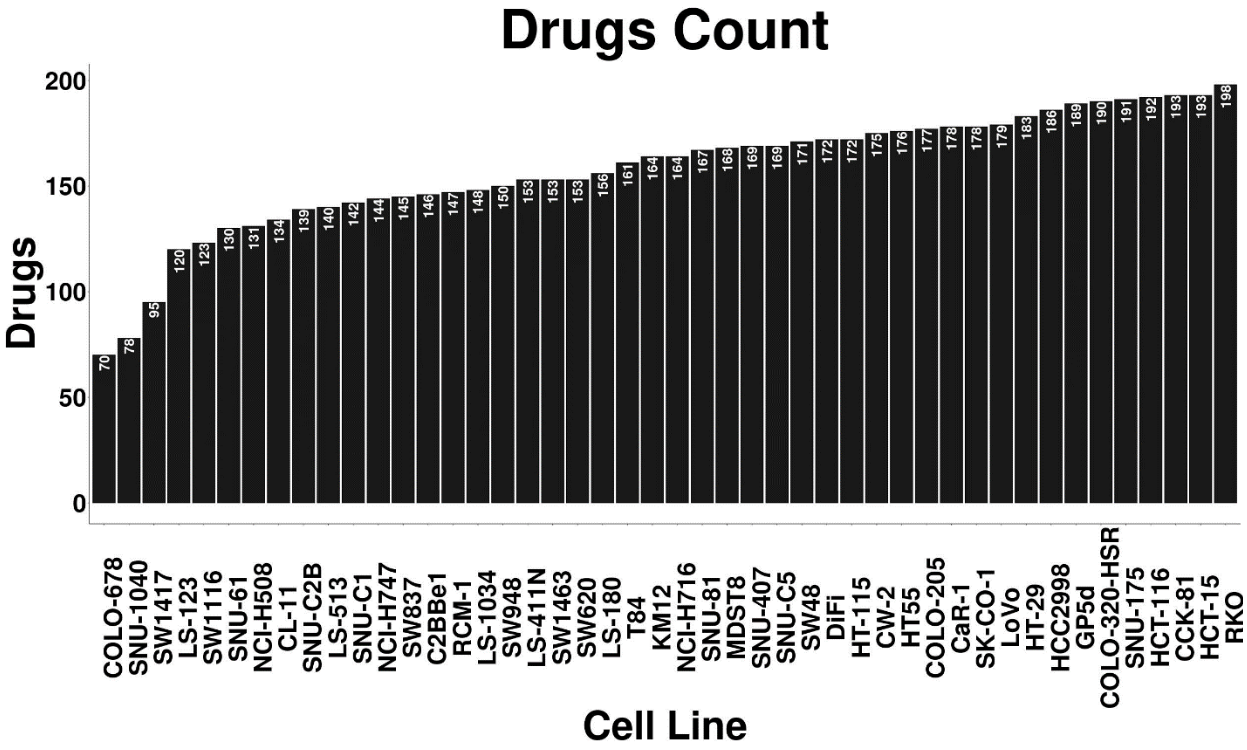

**Supplementary Figure 2:** Scatter plots descripting the relationship between the Actual $logIC_{50}$ values and the predicted $logIC_{50}$ values by our developed QSAR models across the 12 cell lines

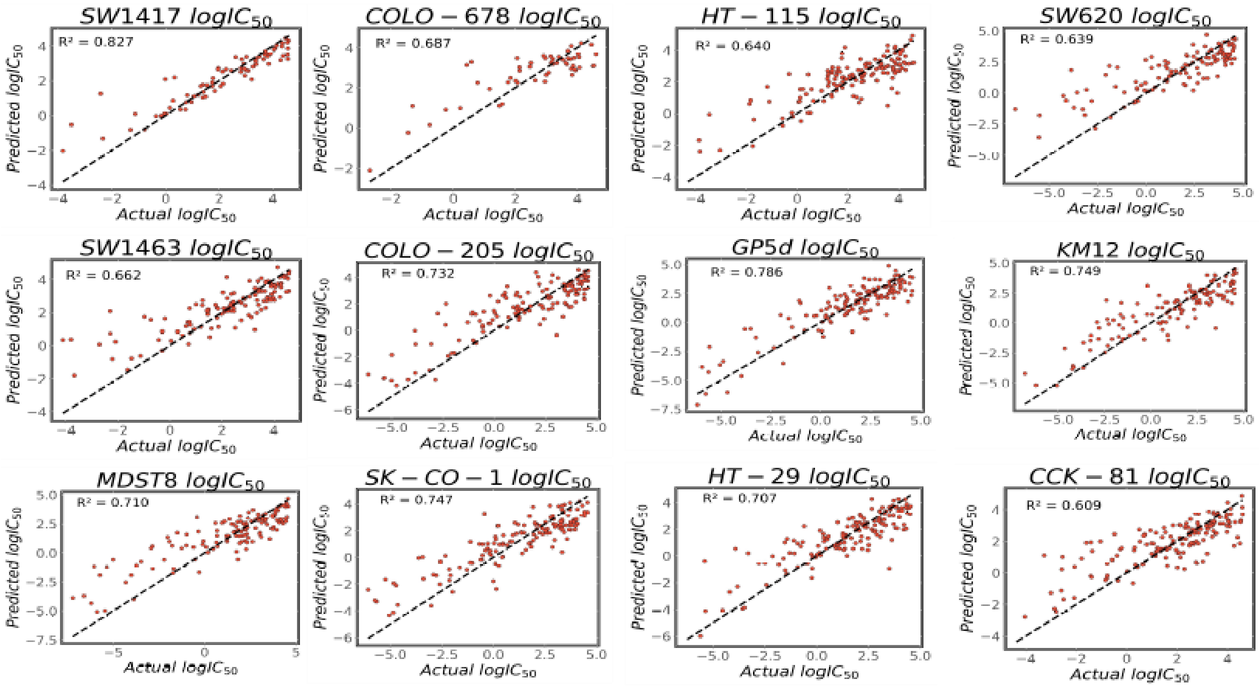

**Supplementary Figure 3: Graphical representation of drug-oncogene relationships of various drug-gene combinations**. To validate our models, we performed the drug-to-oncogene relationship analyses. Our analyses, shows that the A) Muations in *ABCA13* reduced the sensitivity of CRC cell lines for the drug SCH772984, B) Mutations in *ABCA13* increased the sensitivity of CRC cell lines for the drug dinaciclib, C) Mutations in *DNAH9* increased the sensitivity of CRC cell lines for the drug sabutoclax, D) Mutations in *DNAH9* increased the sensitivity of CRC cell lines for the drug vincristine, E) Mutations in *DNAH6* increased the sensitivity of CRC cell lines for the drug SGC0946, F) Mutations in *FLG* increased the sensitivity of CRC cell lines for the drug podophyllotoxin bromide

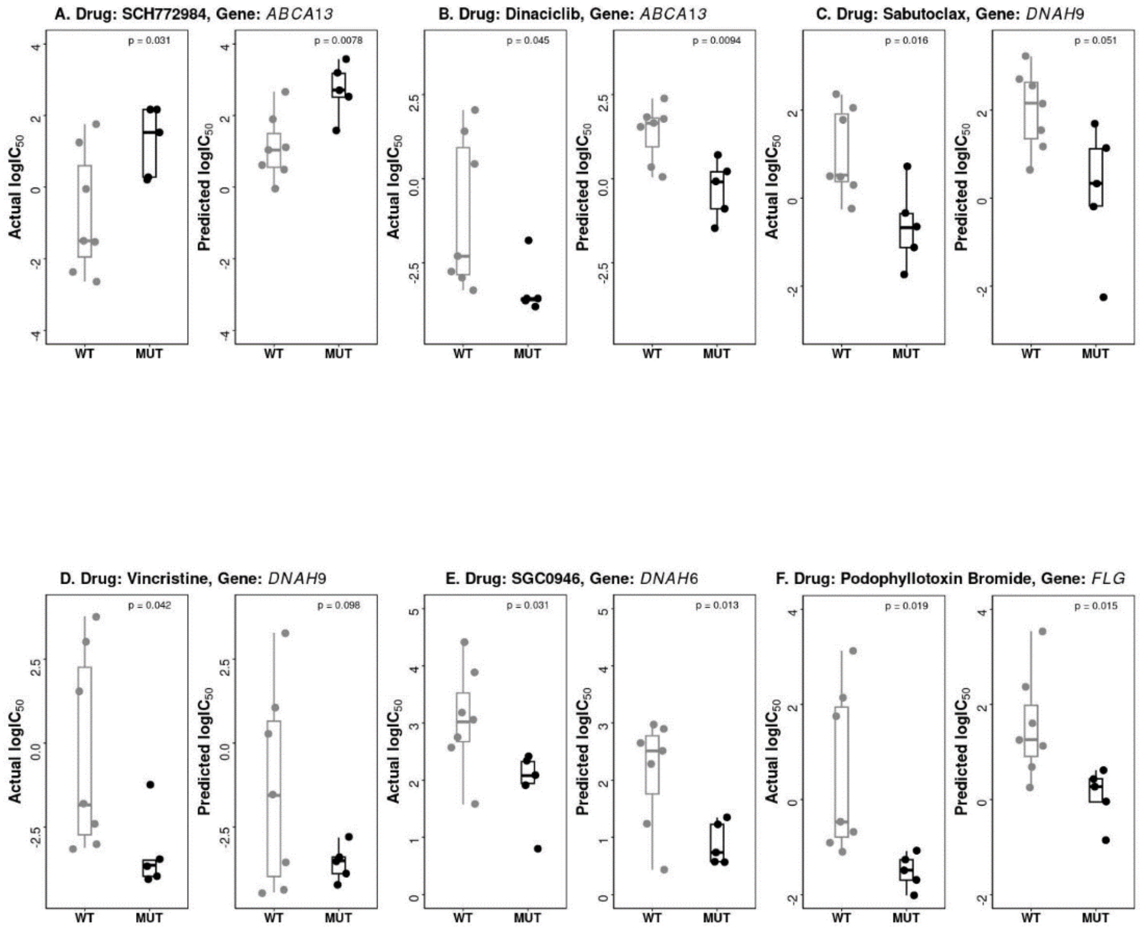
